# Functional inertia reveals history-dependent organization of large-scale brain dynamics

**DOI:** 10.1101/2025.09.30.679686

**Authors:** Sir-Lord Wiafe, Vince D. Calhoun

## Abstract

The brain does not process the present in isolation. Its responses are shaped by the accumulating weight of prior states, yet existing frameworks treat large-scale brain dynamics as sequences of transient configurations, leaving the constraining influence of history implicit in how models are specified rather than measured directly. Here we define functional inertia as the rate of change of an accumulated latent state and make it measurable through an inertial state-space model. Applied to resting-state fMRI in 160 healthy controls and 151 individuals with schizophrenia, functional inertia organizes brain activity into three recurrent regimes, and patients redistribute their time across them in a way that tracks negative symptom severity. The same measure carries opposite consequences in the two groups, linked to better cognition in controls and to more severe symptoms in patients, because what inertia means depends on the accumulated configuration it operates from. The same contextual logic holds across circuits, where stronger inertia tracks greater symptom negative severity in integrative hubs and lower severity in sensory pathways, resolving the longstanding tension between reports of excessive rigidity and reports of excessive volatility in schizophrenia. These findings position functional inertia as a multilevel constraint on brain reorganization, and the accumulated past as a dimension of brain function that can be measured rather than assumed.

## 1. INTRODUCTION

Large-scale brain dynamics are typically modeled as sequences of transient configurations, implicitly assuming that present activity is governed by instantaneous states^1–4^. Yet this assumption is incomplete: identical inputs can produce divergent neural responses not because the inputs differ but because the history of the system does^5–8^. Recurrent excitation^9,10^, NMDA receptor dynamics^9^, synaptic facilitation^11,12^, and neuromodulatory gain control^13–15^ together ensure that prior states persist and constrain how present activity evolves, yet none of these frameworks estimate the accumulated past itself as a quantity that can be measured and compared across individuals. From dynamic functional connectivity and hidden Markov models^1–4^ to whole-brain generative models^16^ and predictive coding, activity is treated as sequences of transient configurations or learned distributional priors. Dynamic causal modelling^17^ and hidden Markov models with persistence structure encode dependence on the past through state equations and transition priors, respectively. However, history enters as a parameter governing how states succeed one another rather than as a quantity carried forward and measured. The constraining influence of history on present reorganization remains implicit in how these models are specified rather than measured directly.

This gap is not merely technical. The brain does not simply transition between states: it evolves under accumulated constraints imposed by prior activity^5–8^. Microcircuit models track moment-to-moment updates while leaving cumulative trajectories implicit^9,10^. Mesoscopic and electrophysiological approaches characterize evolving patterns but analyze state evolution relative to instantaneous configurations rather than accumulated context^6,18^. Even predictive coding and Bayesian brain theories, which most explicitly engage with the role of prior experience, formalize history as learned distributional priors rather than as a cumulative constraint on ongoing state evolution^19–21^. Forecast-based approaches, which quantify how predictable cortical dynamics are from their own recent past using linear forecasting and dynamic mode decomposition^22^, measure how well the next state can be anticipated rather than how much the accumulated state resists being changed. What is missing is not a better description of brain states but a way of measuring the accumulated state itself, so that the degree to which it resists reorganization becomes an observable quantity rather than a modelling assumption.

Here we make that quantity measurable. We define functional inertia as the degree to which accumulated prior states resist ongoing neural reorganization and develop an inertial state-space model that treats accumulated context itself as a state variable, recursively integrating observations into a running latent trajectory whose evolution reflects the cumulative influence of prior states. Functional inertia is operationalized as the rate at which this accumulated trajectory changes, providing a quantitative measure of how strongly history constrains present reorganization. Crucially, the framework is agnostic to the form of observation and can be instantiated using spatial activity maps, regional or multivariate time series, dynamic connectivity trajectories, behavioral sequences, neuromodulatory signals, or any temporally evolving biological observable. Different observational axes provide alternative projections of the same underlying constraint rather than distinct mechanisms. The model is therefore well posed on any such observation, although the findings it returns will depend on which one is chosen.

We instantiate this framework using our recently introduced time-resolved dynamic time warping approach^23^ applied to resting-state fMRI, leveraging its natural cumulative alignment structure^24^ consistent with the temporal integration central to the ISSM. What is new here is the accumulation step and everything built on it, from the latent trajectory to its rate of change, functional inertia, and the ordering between them. Functional inertia emerges as a coherent organizing constraint expressed consistently across three complementary levels of brain organization. At the dynamical level, it structures whole-brain activity into recurrent regimes reflecting distinct modes of history-constrained state evolution. At the whole-brain level, a stable inertial magnitude links these dynamics to cognitive performance and negative symptom severity. At the circuit level, functional inertia is distributed across specific interregional pathways in patterns that differentiate normative and pathological dynamics. We use schizophrenia as a perturbative test case, a disorder with well-documented disruptions in brain dynamics whose mechanistic basis remains contested. Departures from normative inertial organization emerge at all three levels, and they are not independent of one another, since where the accumulated state has settled proves to be upstream of how fast it continues to change.

Ablation analyses further establish the necessity of cumulative integration. Removing the accumulation step abolishes system-level inertial magnitude associations and symptom effects while preserving all other components of the model, demonstrating that accumulated context is not a redundant preprocessing step but the specific operation through which local regime dynamics cohere into a global organizing constraint. A similar three-regime architecture and group differences arise when sliding window Pearson correlation^1^ or phase synchrony^25^ is accumulated instead, so the model is not specific to the choice of observation, although the specific findings and interpretations are. Together these findings establish functional inertia as a measurable property of brain dynamics, informative about both cognition and clinical phenotype, and open a new axis along which healthy and disordered brain function can be studied.

## 2. METHODS

### 2.1. Inertial state-space formulation of brain dynamics

To formalize the idea that human cognition is shaped by context-dependent accumulation of experience, we adopt an inertial state-space model (ISSM). At an abstract level, this describes how a person’s momentary internal state reflects a weighted history of prior states, with past configurations persisting and being differentially expressed depending on the current context. Here, we operationalize this constraint using brain data, treating large-scale neural activity as an observable proxy for the underlying cognitive state illustrated schematically in Fig. 1c. The formulation is observational, providing a compact mathematical description of how observations across time contribute to the present latent configuration.

**Fig. 1:**
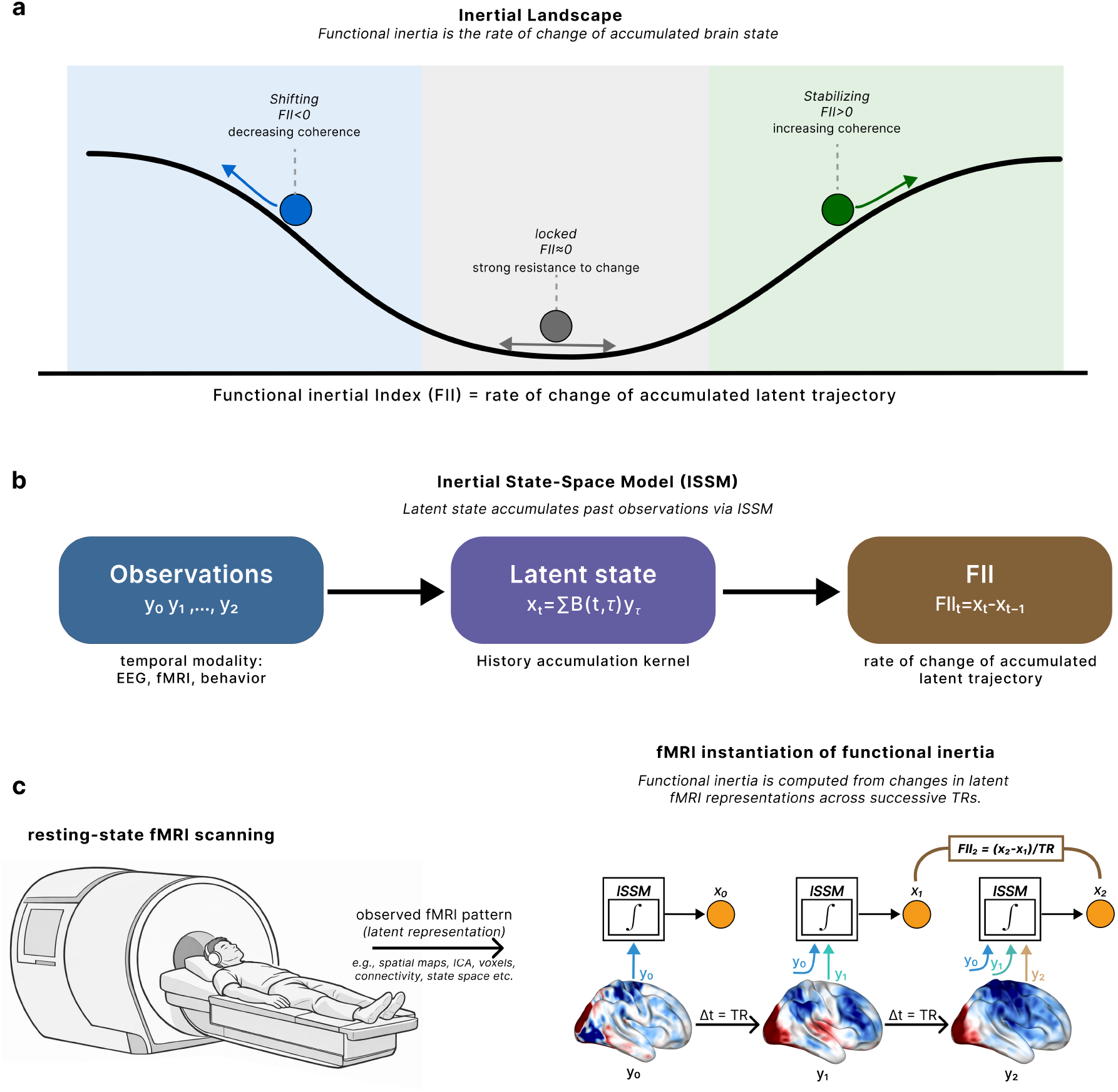
Functional inertia as a history-dependent constraint on brain state evolution. **(a)** Inertial landscape of functional inertia. Functional inertia is the rate of change of the accumulated brain state. The inertial landscape illustrates three recurrent dynamical regimes defined by the sign and magnitude of the functional inertia index (FII). In the locked regime (FII ≈ 0), the system exhibits strong resistance to reconfiguration. In the shifting regime (FII < 0), the system actively reconfigures toward decreasing coherence. In the stabilizing regime (FII > 0), the system actively reconfigures toward increasing coherence. **(b)** Inertial state-space model (ISSM). The latent state x_t_ is defined as a context-modulated accumulation of past observations y_τ_ through an accumulation kernel B(t, τ). The functional inertia index is operationalized as the rate of change of this accumulated latent trajectory (FII_t_ = x_t_ − x_t–1_). The formulation is representation-agnostic and can be instantiated across neural, behavioral, or physiological observables. **(c)** fMRI instantiation of functional inertia. fMRI observations are mapped to latent representations across successive repetition times via the ISSM. Each successive integration step accumulates prior observations, and changes in the accumulated latent trajectory are used to compute the functional inertia index at each time point.

Let *x*(*t*) denote the latent state of the system at time *t*, corresponding to the internal cognitive configuration depicted in the schematic model, and let *y*(*t*) denote the observed multivariate brain signal used to infer this state. In its most general continuous-time form, the inertial state-space model is defined as

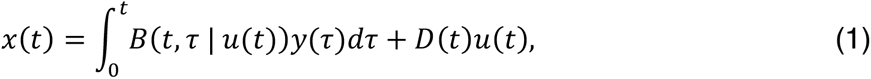

where *B*(*t*, *τ* ∣ *u*(*t*)) is a context-modulated accumulation kernel specifying how prior observations *y*(*τ*) at time indices *τ* ≤ *t* contribute to the current state, and *u*(*t*) represents external inputs or contextual influences when present. The term *D*(*t*)*u*(*t*) captures direct effects of such inputs on the latent state.

Within this formulation, the accumulation kernel defines how present brain states are constructed from observations across time in a context-dependent manner. This captures how state evolution unfolds within a latent space shaped jointly by accumulated history and current context, with functional inertia defined as the rate of change of that accumulated latent state.

For discretely sampled neuroimaging data, such as fMRI, the formulation becomes

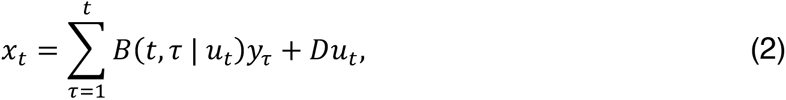

where *x_t_* denotes the latent state at time index *t*, and *y_τ_* the observed signal at time indices *τ* ≤ *t*. An intuitive version of the ISSM formulation is provided in Fig. 1b.

This formulation is deliberately broad. While we focus here on brain-derived observations, the same inertial description applies to human cognition more generally, with different observational modalities, including neural or behavioral readouts, corresponding to different instantiations of *y*(*t*). Different accumulation kernels define alternative operationalizations of inertia, while preserving the constraint that present states reflect the cumulative trajectory leading to the present moment. The empirical work presented here tests only the input-free, zero-order case of this formulation, described next; the general context-modulated form is presented to situate that case.

### 2.2. Zero-order instantiation of the ISSM

The inertial state-space formulation admits a zeroth-order instantiation in which the latent state is defined solely by accumulated observations, without requiring an explicit, parameterized input or task model. This regime arises naturally when state evolution is driven by internally generated dynamics or by contextual influences that are implicit or unmeasured, as in spontaneous thought, mind-wandering, memory recall, affective drift, or naturalistic conditions such as task-free viewing or listening. In these settings, the absence of explicit inputs does not imply a lack of structure; rather, it isolates a principled limit in which the influence of accumulated history on state evolution can be examined directly.

Resting-state fMRI provides a particularly clear and widely studied example of this regime. In the absence of externally imposed tasks, large-scale brain activity reflects internally generated fluctuations shaped by prior configurations of the system. For discretely sampled zero-order ISSM, we set *u_t_* = 0 and equation (2) reduces to

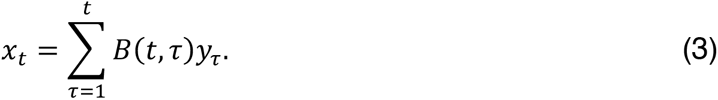

Within this class, the kernel specifies how the influence of past observations is weighted over time. Brain dynamics are shaped more by the recent past than by the distant past, with the influence of a prior configuration fading as time passes^26,27^. The accumulation kernel is where this is expressed, and we therefore adopt an exponentially weighted moving average (EWMA) kernel,

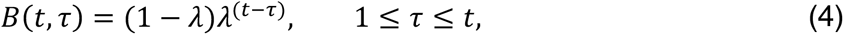

where *λ* ∈ (0, 1) is the retention factor, the proportion of the accumulated state carried forward at each time step, so that an observation *τ* steps in the past is weighted by *λ*^(*t*−*τ*)^ and its influence decays geometrically. Equation (4) yields the ISSM latent representation as the recursion

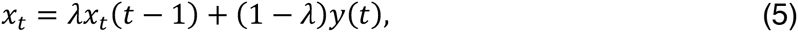

in which the latent state at each time point is a weighted blend of the state accumulated so far and the current observation. The retention factor, *λ*, is set by the memory constant, *τ_m_*, the time in seconds over which influence decays to 1/*e* of its initial value, 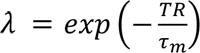, where TR is the repetition time. We set *τ_m_* = 100 *s*, the period corresponding to the 0.01 Hz high-pass applied to the fMRI data^28,29^, so that the latent state integrates over the slowest fluctuations the data retain. This gives *λ* = 0.980 at a TR of 2 s. Sensitivity to this choice is also reported in further analysis.

Every past observation therefore still contributes, but with a weight that falls off with elapsed time, so the latent state is the running configuration implied by the recent trajectory, and functional inertia is revealed by the deviation required for new observations to reorganize it. By isolating state evolution from explicit inputs, the zero-order instantiation exposes inertia as an intrinsic property of brain dynamics while making no assumptions about task structure or contextual drives, a choice that aligns naturally with resting-state conditions in which the relevant influences are endogenous and unmeasured.

### 2.3. Functional Inertia Index (FII) and Functional Inertia Index Magnitude (FIIM)

Within the inertial state-space formulation, functional inertia corresponds to resistance to change in the latent state over time. We operationalize it by quantifying the temporal rate of change of the latent trajectory, which we term the functional inertia index (FII). Because inertia reflects resistance to change, values of FII close to zero indicate strong inertia, whereas larger-magnitude values indicate active reconfiguration. Formally, the FII at time *t* is defined as the temporal derivative of the latent state,

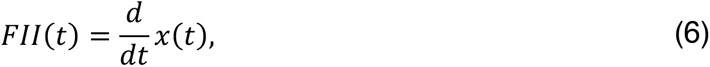

or, in the discrete-time setting,

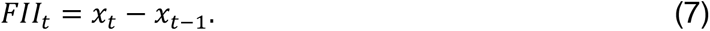

This definition holds regardless of how the latent state is constructed and therefore applies across different instantiations of the inertial state-space model.

The FII is signed, and its sign carries directional information about state evolution: depending on the representational axis, positive values indicate evolution toward a more coherent configuration and negative values evolution toward a more divergent one. We define the absolute trace of the FII as the functional inertia index magnitude (FIIM), which quantifies the magnitude of the reconfiguration. Both FII and FIIM are defined independently of how the latent state is constructed and therefore apply across different instantiations of the inertial state-space model. This quantification assumes no task structure or mechanistic force and provides a time-aware quantification of how much past configurations shape present evolution.

### 2.4. fMRI data & processing

We analyze resting-state fMRI from the Function Biomedical Informatics Research Network (fBIRN) consortium. Data was acquired with a repetition time of 2 s and comprised 160 healthy controls (37.0 ± 10.9 yr, 45 F/115 M) and 151 individuals with schizophrenia (38.8 ± 11.6 yr, 36 F/115 M). Volumes underwent standard preprocessing such as slice-timing correction, rigid-body realignment, MNI spatial normalization, and 6 mm FWHM Gaussian smoothing^30^. Spatially independent components were then extracted using the NeuroMark ICA pipeline^31^, yielding 53 intrinsic connectivity networks (ICNs) common to all subjects. The resulting ICN time courses are detrended, despiked, and filtered using a Butterworth bandpass filter with a frequency cutoff of 0.01 − 0.15*Hz*^32^. The filter, with an optimal order of 7 as determined by MATLAB’s *buttord* function, maintained a passband ripple under 3 *dB* and a minimum stopband attenuation of 30 *dB*. Finally, we standardize the filtered ICNs through z-score transformation.

### 2.5. Dynamic time warping instantiation of the ISSM

The inertial state-space formulation is agnostic to the representational axis used to define the observed trajectory *y*(*t*). Here we instantiate it on time-resolved interregional profiles, defining *y*(*t*) as the dynamic time warping (DTW) distance between the time courses of a pair of intrinsic connectivity networks^23,33,34^. DTW is suited to this role for two reasons: it accommodates the nonuniform timescales of brain dynamics through time-adaptive alignment, and it returns a non-negative, additive distance that accumulates as a deformation cost across aligned time points^24^, matching the algebraic structure of the ISSM. The quantity accumulated is therefore interregional amplitude disparity, and the analysis yields one latent trajectory per pair, or 1,378 trajectories per participant across the 53 ICNs. For each pair, DTW was computed between the two z-scored time courses using a Sakoe–Chiba band^24^ of half-width 50 time points (100 s), matching the memory of the ISSM used here and consistent with the window adopted for DTW in fMRI^33–35^.

The local cost was |*x*(*i*) − *y*(*j*)|, with *γ* = 1.5, a value selected in prior work for test– retest reliability and clinical sensitivity^23^. The optimal warping path was obtained by backtracking, and the local cost at each step along that path was taken as the time-resolved DTW distance^23^. This was resampled to integer time indices by piecewise cubic Hermite interpolation and taken as y(t). Before warping, each time course was reflection-padded by 30 frames at both ends and the padded region discarded afterwards, so that the alignment at the start and end of the scan is not biased by the series boundary. The latent state *x*(*t*) was then accumulated from *y*(*t*) using equation (5). Because direct differencing amplifies high-frequency noise, the latent state was smoothed with a Gaussian kernel of FWHM 6 frames before differencing, with the bandwidth set from the elbow of the subject-averaged power spectral density of the latent trajectories (Supplementary Fig. 1). The full derivation, covering the local cost, the cumulative cost recursion, and the time-resolved distance taken along the warping path, is given in the Supplementary Methods.

The FII was then taken as the first difference of the smoothed trajectory and multiplied by −1, so that positive FII corresponds to reconfiguration toward reduced interregional disparity and negative FII to reconfiguration toward increased disparity. Averaging the absolute value of the FII, FIIM, over time gives one **edge-level FIIM** per brain network pair; averaging that across the 1,378 pairs gives the participant’s **whole-brain FIIM**. The accumulated latent state is summarized the same way, giving an edge-level and a whole-brain accumulated latent state.

#### 2.5.1. Anchor baseline window

Cumulative estimates from the earliest frames are unstable, so each scan is given an anchor window that establishes the reference configuration. After discarding the first 10 s for magnetization equilibrium^36^, the anchor comprises the following 50 time points (100 s) to give room for early time points to realize the EWMA memory chosen. The latent state is initialized on the mean of this window, so FII begins at frame 51 and 157 volumes yield 107 FII time points. The anchor is a within-subject reference, not an assumed common state across individuals. All 107 FII frames enter every analysis; boundary effects are handled by the reflection padding rather than by discarding frames.

### 2.6. Recurrent functional inertia regime estimation

At each time point, the FII across all 1,378 region pairs forms a vector describing the whole-brain pattern of reconfiguration at that moment. These vectors are clustered with k-means, which is standard in time-resolved fMRI to yield distinct temporal regimes of the FII^37^. Time points across subjects are concatenated into a single time-by-feature matrix (time points as samples, edges as features) prior to k-means clustering. We implement the *k*-means algorithm with cluster numbers ranging from 1 to 10, setting the maximum number of iterations to 10,000 and using 20 random initializations to ensure robust convergence. Each centroid is therefore a vector of 1,378 values, one per region pair. The city-block (Manhattan) distance metric is selected for its demonstrated robustness in handling high-dimensional data^38^. The optimal cluster number is determined as 3 using the elbow criterion (See Supplementary Figure 2).

### 2.7. Temporal regime quantification

Using *k*-means clustering, we characterized each participant’s temporal behavior across the estimated temporal regimes with three metrics. First, the mean dwell time (MDT) measures regime persistence by averaging how long a participant remains in a state once it is entered^39^. Second, the fraction rate (FR) reflects overall occupancy by expressing the proportion of the total recording spent in each state^39^. Finally, the regime-to-regime transition probabilities form the transition matrix (TM), which quantifies the likelihood of a participant moving from one regime to any other^39^. In this study, persistence refers to MDT and regime engagement to FR and MDT jointly. The FII and the accumulated latent state are also summarized within each regime by computing the temporal average for time points assigned to it.

### 2.8. Group analysis & regression models

Group differences between healthy controls and individuals with schizophrenia, associations with Computerized Multiphasic Interactive Neurocognitive System (CMINDS) cognitive scores, and associations with Positive and Negative Syndrome Scale (PANSS) symptom scores are assessed using generalized linear models including age, sex, scanning site, and mean framewise displacement (FD) as covariates. Group differences use all participants; symptom associations are estimated within the schizophrenia group, and cognitive associations within each group separately. A Gaussian family is used for group differences and CMINDS scores, and a Poisson family for PANSS scores, which are non-negative integer counts with a right-skewed distribution ^23,40^. Effect sizes are reported as standardized coefficients, in units of outcome change per standard deviation of the predictor. Multiple comparisons are controlled with the Benjamini-Hochberg false discovery rate procedure, applied within each metric family: across the three regimes for FR, across the three for MDT, across the nine entries of the TM, and across the 1,378 region pairs for edge-level analyses. Whole-brain FIIM is tested once per outcome, so no correction applies. Joint distributions of the latent state and the FII are shown as two-dimensional kernel density estimates pooled over participants and frames, with contours drawn at fixed density levels. Where indicated, regime effects are refitted with whole-brain accumulated latent state or medication added as covariates, to test whether they are separable from the position of the latent state or medication effects respectively.

### 2.9. Comparison with conventional dynamic connectivity

To test whether the FII time course restates conventional dynamic functional network connectivity (dFNC), the two are compared directly at every region pair. For each participant and pair, sliding window Pearson correlation is computed over windows of 22, 44, 88 and 100 s, stepped one frame at a time and centered on the same frames as the FII, giving one conventional time course per window length. Each is then correlated with the FII time course at that pair, and the squared correlation is taken as the variance the two signals share. The instantaneous DTW distance *y*(*t)* is included as a reference, being the signal the FII is built from. Because both observations are heavily autocorrelated, the sampling distribution of this correlation is much wider than its nominal degrees of freedom imply. We therefore build a null by circularly shifting one of the two signals within a participant, which preserves each signal’s own autocorrelation exactly and destroys only the temporal correspondence between them, repeated over 20 shifts per pair. Shared variance is reported against the shared variance expected under this null, so a pair lying on the identity line shares no more than chance.

### 2.10. Ordering of the latent state and FIIM

Alongside the FII, the accumulated latent state differs between individuals. For each participant, we average it over time and across the 1,378 region pairs, giving a whole-brain accumulated latent state, the counterpart of whole-brain FIIM. To establish which of the two is upstream, we ask whether the association of diagnosis with whole-brain FIIM survives the inclusion of the latent state, and whether the association of diagnosis with the latent state survives the inclusion of whole-brain FIIM. The two are then entered together as predictors of each PANSS score, both for the whole scan and restricted to the frames assigned to each regime in turn. Edge-level association maps are compared between analyses by Pearson correlation across the 1,378 pairs.

### 2.11. Alternative observation models

The ISSM is agnostic to the choice of *y*(*t*), so the DTW instantiation is compared against two others derived from the same preprocessed ICN time courses: sliding window Pearson correlation^1^ and phase synchrony^25,28^. For sliding window correlation, Pearson correlation is computed between each pair over a 22-timepoint window with a step size of 1 timepoint, and *y*(*t*) is taken as 1 − *r*(*t*). For phase synchrony, the instantaneous phase of each time course is obtained by wavelet transform^41,42^, and *y*(*t*) is taken as 1 − *cos*(*φ*ᵢ − *φⱼ*). Both are disparity-like, in the same direction as DTW distance, so larger values mean the two regions are further apart.

Each observation is then put through the identical pipeline: the same EWMA kernel, the same anchor window, the same smoothing, differencing, and clustering. The regimes that emerge are not directly comparable across observation models, because only the DTW regimes are single signed; regimes are therefore ranked low, mid, and high within each model by the median absolute value of their centroid and compared on that ordering.

### 2.12. Robustness analyses

#### 2.12.1. Accumulation and memory length

The kernel is varied while everything else is held fixed. In the ablation, accumulation is removed entirely, and the FII is taken by smoothing and differencing *y*(*t)* directly, so the latent state plays no part. The EWMA kernel is then applied at memory constants of 30, 60, 100, and 120 s, and finally replaced by the uniform cumulative kernel, which weights the whole scan equally. Whole-brain and edge-level analyses are repeated under each. At the edge level, the sets of region pairs surviving FDR under each kernel are compared against the ablation by McNemar test. The test asks whether the number of pairs gained under the kernel exceeds the number lost relative to the ablation.

#### 2.12.2. Reproducibility across participants

Reproducibility is assessed by splitting the participants at random into two halves, stratified by diagnosis and scanning site; the mean edge-level FIIM map is computed in each half, and the two maps are correlated across the 1,378 pairs. This is repeated over 200 random splits.

#### 2.12.3. Elapsed scan time

To test whether the index decays as the scan proceeds, mean FIIM is binned into quarters of the scan and expressed relative to each observation model’s own scan average, and a power-law exponent is fitted to its dependence on elapsed time.

#### 2.12.4. Head motion

Patients move more than controls, so the group difference in whole-brain FIIM is refitted under progressively stricter exclusion on mean framewise displacement, at thresholds of 0.40, 0.30, 0.25 and 0.20 mm alongside the full sample. Framewise displacement remains a covariate in every model.

#### 2.12.5. Medication

Chlorpromazine-equivalent dose is available for 99 of the 125 patients, so it is examined as a sensitivity analysis rather than a standard covariate. The negative-symptom regime effects are refitted with dose added and compared against the same models fitted on the identical subsample without it, so that any change is attributable to the covariate rather than to the smaller sample.

## 3. RESULTS

### 3.1. Recurrent functional inertia states reflect whole-brain dynamical regimes

We instantiated the ISSM using dynamic time warping applied to interregional fMRI time series and defined the functional inertia index as the rate of change of the resulting accumulated latent trajectory, such that values near zero reflect strong resistance to reconfiguration and larger absolute values reflect active reconfiguration away from or toward a more cohesive amplitude configuration (Fig. 2a). To test whether FII exhibits structured dynamical organization, we applied k-means clustering to FII values from all interregional pairs and time points concatenated across subjects. The resulting regimes exhibited strong whole-brain segregation (Fig. 2b). Two of them are single-signed: in one, every one of the 1,378 region pairs has positive FII, spanning +3.9 to +12.4 × 10⁻⁴, so the whole brain is moving toward reduced interregional disparity, and we term this the **stabilizing** regime. In the other, every pair has negative FII, spanning −14.8 to −4.1 × 10⁻⁴, so disparity is growing everywhere, and we term this the **shifting** regime. The third sits at zero and straddles it, reflecting moments when the accumulated configuration barely changes, and we term this the **locked** regime. These are therefore global modes of history-dependent evolution rather than distinct network configurations.

**Fig. 2:**
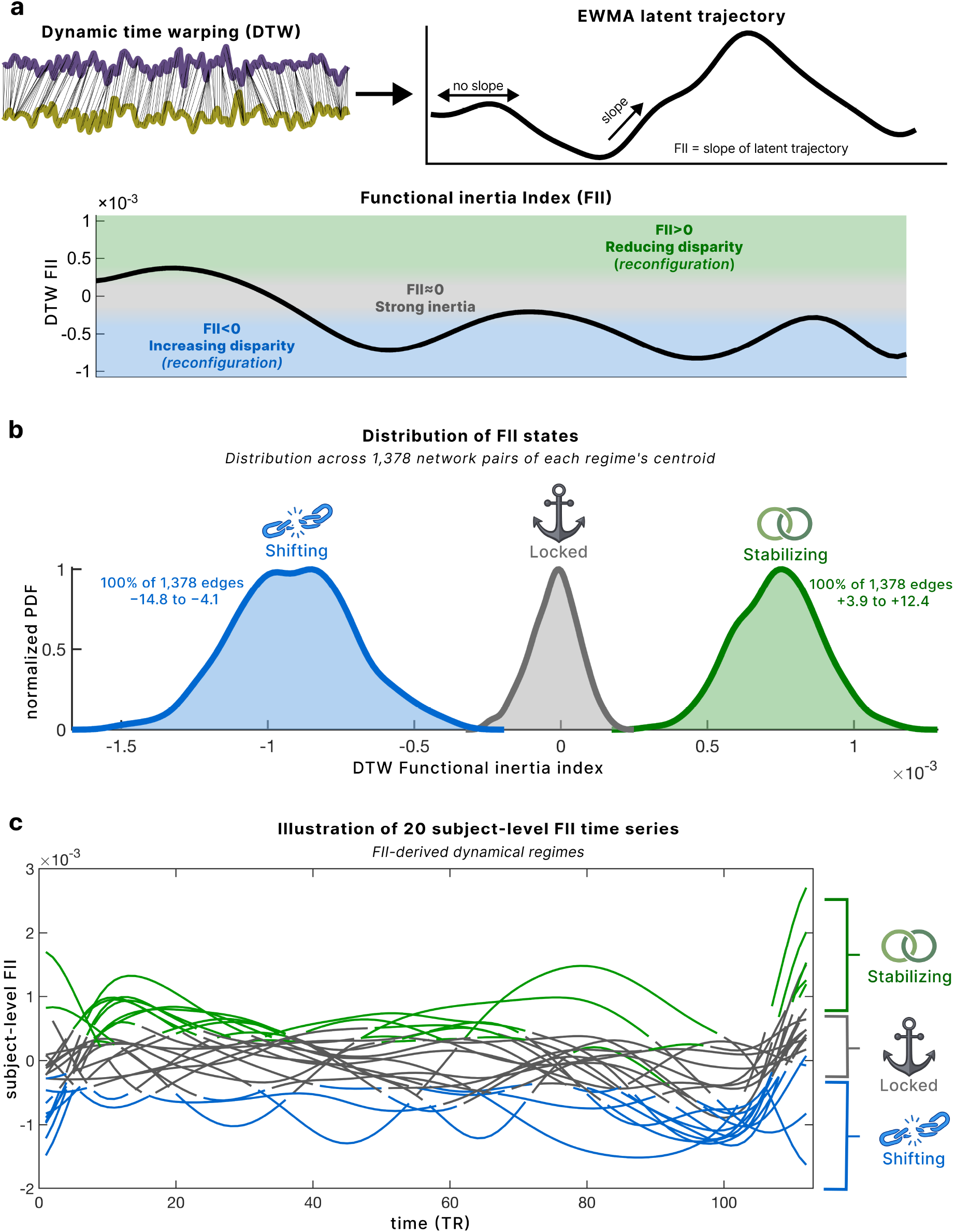
Dynamic time warping instantiation of functional inertia and emergent dynamical regimes. **(a)** Dynamic time warping (DTW) aligns interregional amplitude trajectories while accommodating intrinsic timescale variability, yielding a time-resolved deviation signal. This deviation is recursively accumulated with an exponentially weighted moving average (EWMA) to form a latent trajectory that reflects the integrated history of prior observations. The functional inertia index (FII) is defined as the temporal rate of change of this accumulated trajectory. The functional inertia index is defined as the temporal rate of change of this accumulated trajectory, with values near zero reflecting strong resistance to reconfiguration and nonzero values reflecting active reorganization. **(b)** Distribution of functional inertia states across the whole brain. Pooling FII values across all subjects, time points, and interregional pairs reveals a trimodal distribution corresponding to three distinct dynamical regimes. These regimes are centered around all negative FII values (shifting), near-zero FII values (locked), and all positive FII values (stabilizing), indicating that functional inertia organizes brain dynamics into qualitatively distinct modes of state evolution. **(c)** Subject-level functional inertia time series illustrating recurrent dynamical regimes. Representative FII trajectories from 20 individuals show that these regimes recur over time within individuals and are expressed coherently across interregional pairs.

We identify three recurrent regimes. The locked regime, characterized by FII values near zero, reflects strong resistance to reconfiguration. The stabilizing regime, with positive FII values, reflects active reconfiguration toward a more cohesive amplitude configuration with decreasing interregional disparity. The shifting regime, with negative FII values, reflects reconfiguration toward a more dispersed configuration with increasing interregional disparity. Individual FII trajectories illustrate how participants move among the three regimes over the course of a scan (Fig. 2c). Functional inertia therefore organizes brain activity into three recurrent whole-brain regimes that differ in both the direction and the amount of history-dependent reconfiguration.

### 3.2. Altered engagement with functional inertia regimes in schizophrenia

Schizophrenia provided a perturbative test of whether functional inertia reflects a genuine organizing constraint: if accumulated history governs large-scale dynamics, disorders characterized by disrupted brain organization should exhibit systematic alterations in how inertial regimes are expressed and engaged.

The two groups (patients and controls) differ first in where they sit. Plotting each participant’s accumulated latent state against their FII places every moment of every scan in a plane whose horizontal axis is how far the accumulated configuration has drifted and whose vertical axis is how fast it is changing (Fig. 3a). The patient cloud is displaced to the right of the control cloud across the whole vertical range, so patients carry greater accumulated latent interregional disparity whether they are reconfiguring quickly or barely at all. Controls spend 56% of the scan in the locked regime against 46% in patients, with the difference redistributed into stabilizing (21% versus 30%). This latent displacement is near-global at the edge level, with 891 of the 1,378 region pairs elevated in patients and none reduced (minimum *q* = 3.9 × 10⁻⁶; Fig. 3b), and it holds separately within each of the three inertial regimes, including the locked one (all *q* < 0.001; Fig. 3c). It is therefore not a by-product of the two groups occupying different regimes.

**Fig. 3:**
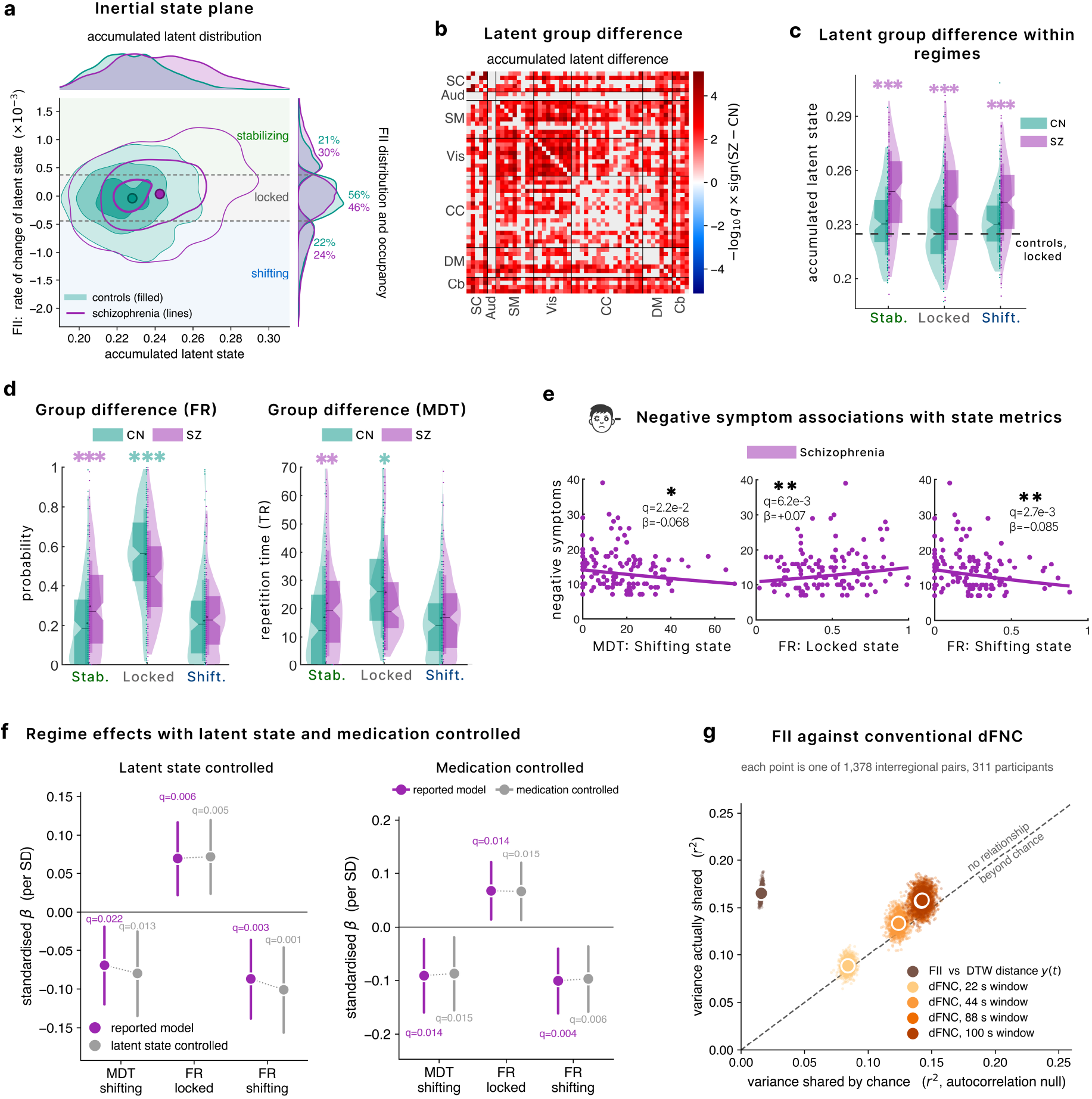
Regime structure of functional inertia and its relation to diagnosis and negative symptoms. **(a)** Joint distribution of the accumulated latent state (x, horizontal) and the functional inertia index (FII = −dx/dt, vertical) across all participants and time points, for controls (teal, filled) and schizophrenia (purple, outlines). The FII distribution on the right shows the three k-means regimes: stabilizing (FII > 0), locked (FII ≈ 0), and shifting (FII < 0), with each group’s occupancy beside it. Controls occupy the locked regime 56% of the scan against 47% in patients, with the difference redistributed into stabilizing (22% vs 29%). Contours are kernel-density levels pooled over participants and frames; at corresponding density levels, patient contours sit right of control contours at every level and across the whole FII range. **(b)** Edgewise group difference in accumulated latent magnitude across the 1,378 interregional pairs; grey indicates pairs not surviving BH-FDR. 891 of 1,378 pairs are elevated in patients, and none are reduced. **(c)** Accumulated latent state within each regime by group. Patients sit above controls in all three regimes (all q < 0.001). **(d)** Group differences in fraction rate (FR, left) and mean dwell time (MDT, right) for each regime, covariate-adjusted for age, sex, site, and framewise displacement, FDR-corrected within each metric family. Patients show higher stabilizing and lower locked occupancy. **(e)** Within-patient associations between regime metrics and PANSS negative symptom score (Poisson, same covariates, n = 125): MDT shifting (q = 0.022), FR locked (q = 0.0062), and FR shifting (q = 0.0027). More time in the locked regime is associated with higher negative symptoms, whereas more time in the shifting regime is associated with lower scores. **(f)** The same three effects refitted with whole-brain accumulated latent magnitude (left panel) and medication (right panel) added as a covariate (grey) beside the reported model (purple). All three survive, so the regime metrics are not a re-description of the latent space where the system sits. **(h)** Variance shared between the FII time course and conventional dynamic connectivity, against the variance expected under circular time-shifts null to preserve the autocorrelation; each point is one of 1,378 pairs. Against sliding-window dFNC, the variance shared falls on the identity line at every window length, meaning it shares no more variance than chance. The DTW distance the FII is derived from shows small overlap (r² ≈ 0.17). Therefore, FII is not a restatement of dFNC or the DTW distance. Throughout, teal is controls and purple is schizophrenia; ***q < 0.001, **q < 0.01, *q < 0.05.

Inertial regime engagement through fraction rate and dwell time also differs (Fig. 3d). Patients spent more of the scan in the stabilizing regime and remained there longer once entered, indexed by fraction rate and mean dwell time; controls showed the inverse in the locked regime. The shifting regime did not differ, and no transition probability survived correction. Within the schizophrenia group, regime engagement tracked negative symptom severity (Fig. 3e). Higher negative symptom severity was associated with greater occupancy of the locked regime and with reduced engagement of the shifting regime, indexed by lower FR and shorter dwell time. These symptom effects are not a restatement of where the system sits, nor an artefact of medication (Fig. 3f). Refitting each with whole-brain accumulated latent state as an additional covariate left all three intact and slightly strengthened. Refitting them with chlorpromazine-equivalent dose added, on the 99 patients for whom dose was recorded, changed the coefficients by less than 0.01 and left all three significant.

Finally, we show that the FII time course is not a relabeling of conventional dynamic connectivity (Fig. 3g). Compared with sliding window Pearson correlation dFNC at windows from 22 to 100 s, it shares no more variance than expected under a circular shift null, which breaks their correspondence while leaving each signal’s own temporal structure intact. The only comparison that exceeds chance is with the DTW distance from which the FII is derived, and even there the shared variance is 14% against 1% expected. Our findings show that schizophrenia is not characterized by uniform disruption of inertial dynamics. The accumulated state is displaced almost everywhere, while the redistribution of time across inertial regimes is selective, and it is that redistribution which tracks negative symptom severity.

### 3.3. Whole-brain and circuit-level inertia dissociate cognition from symptoms

Regime engagement describes how the inertial constraint is expressed moment to moment. Averaged over the whole scan, it also yields a single number per participant, the whole-brain FIIM, where lower values mean the accumulated configuration changes less and inertia is stronger. This single number carries opposite consequences in the two groups (Fig. 4a). Among patients, lower whole-brain FIIM accompanied greater negative symptom severity (*q* = 0.0026), so those whose brains reconfigured least were the most symptomatic. Among controls, lower FIIM was likewise associated with better performance on the CMINDS composite (*q* = 0.002), reasoning and problem solving (*q* = 0.044), and visual learning (*q* = 0.024). Stronger inertia therefore predicts cognitive advantage in health and symptom burden in schizophrenia.

**Fig. 4:**
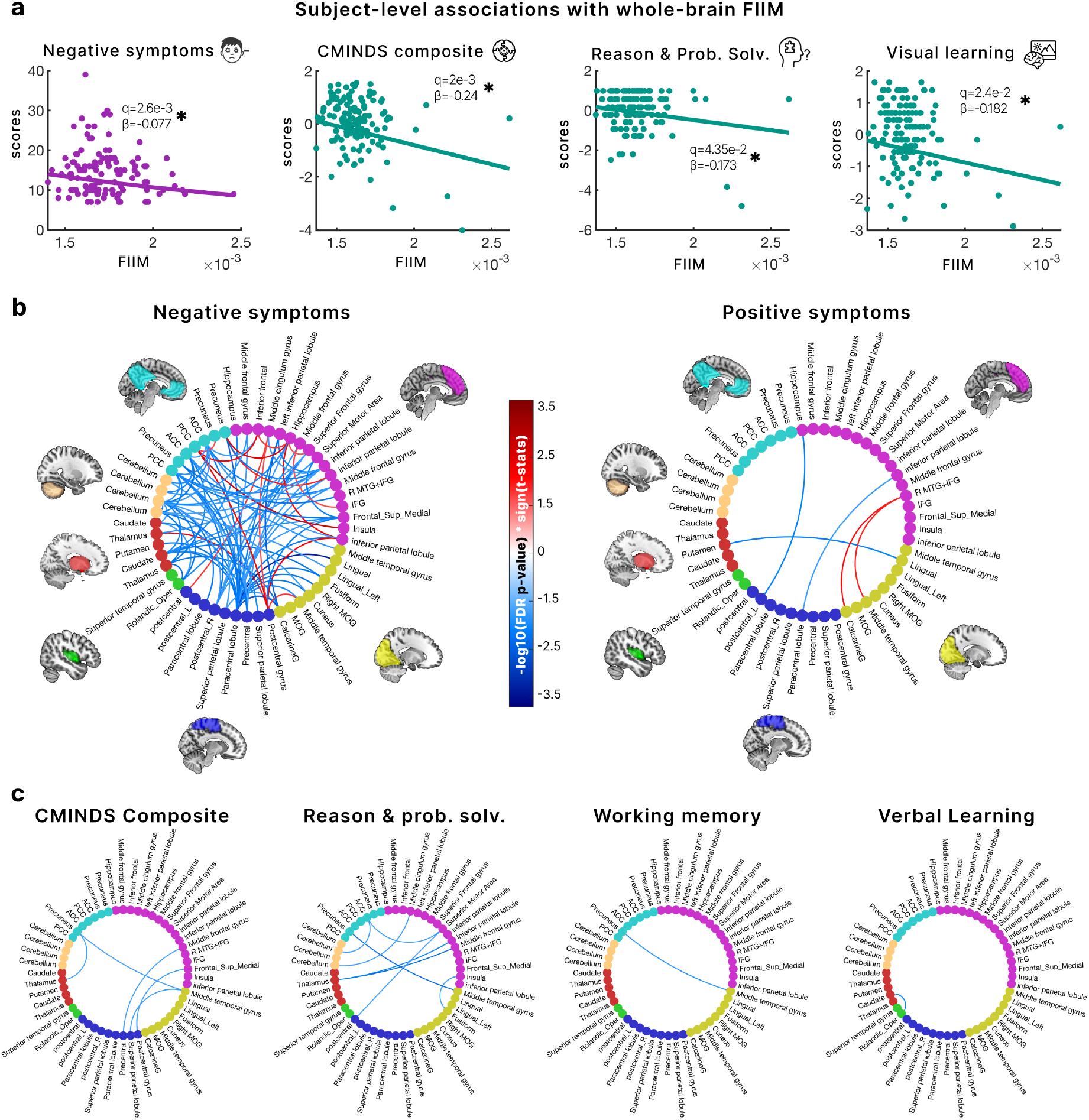
Whole-brain and edge-wise functional inertia index magnitude map onto distinct symptom and cognitive circuits across health and disorder. **(a)** Subject-level associations between whole-brain functional inertia index magnitude (FIIM) and clinical and cognitive measures. Lower FIIM, reflecting stronger overall functional inertia expressed across time and interregional pairs, is associated with greater negative symptom severity in schizophrenia (purple) and with poorer cognitive performance in healthy controls (teal). Each point represents a subject; lines indicate fitted regression relationships after covariate adjustment. **(b)** Edge-wise associations between FIIM and symptom severity in schizophrenia, shown as connectograms over the 53 NeuroMark intrinsic connectivity networks grouped by domain. Edges are colored by signed FDR significance, so blue indicates that stronger inertia at that pair accompanies greater severity and red the reverse. Negative symptoms engage a widely distributed set of cerebellar, subcortical, and default-mode pairs. **(c)** The same analysis for cognitive performance in healthy controls. Fewer pairs survive, and all point the same way: stronger inertia at those pairs accompanies better performance. Together, functional inertia constrains the system in a group-dependent way, and symptoms and cognition draw on largely separate circuits.

Resolving the same measure at each of the 1,378 region pairs shows that these two consequences occupy largely separate circuits. Within patients, 92 pairs were associated with negative symptom severity (Fig. 4b), concentrated in cerebellar, subcortical and default-mode couplings; 77 ran in the direction where stronger inertia accompanies greater severity, and 15 in the reverse. Positive symptoms engaged only 5 pairs. Within controls, far fewer pairs survived, and all ran the same way, stronger inertia accompanying better performance: 8 for reasoning and problem solving, 5 for the composite, and 1 each for working memory and verbal learning (Fig. 4c). Functional inertia is therefore not expressed uniformly across the brain. It acts as a system-level property whose consequences depend on diagnostic context, and the circuits through which it reaches symptoms are largely not the circuits through which it reaches cognition.

### 3.4. The accumulated latent state is upstream of inertial magnitude

The analyses so far describe two quantities that both differ in schizophrenia: where the accumulated configuration sits, and how fast it changes (inertia). Because they are derived from the same trajectory, we asked whether one is upstream of the other (Fig. 5a). Diagnosis predicted the accumulated latent state (*β*/*SD* = +0.61, *P* < 1 × 10⁻⁴) and the latent state predicted whole-brain FIIM (*β*/*SD* = +0.55, *P* = 1 × 10⁻¹⁸). The association between diagnosis and FIIM, which is substantial on its own (*β*/*SD* = +0.44, *P* = 3 × 10⁻⁴), disappeared once the latent state was included (*β*/*SD* = +0.10, *P* = 0.34). The reverse did not hold: diagnosis still predicted the latent state with FIIM in the model (*β*/*SD* = +0.41, *P* < 1 × 10⁻⁴). Only one direction collapses, which establishes the accumulated latent state as upstream of inertial magnitude rather than the two being interchangeable. Patients therefore do not reconfigure faster for some independent reason; the elevated FIIM in schizophrenia is downstream of their displaced accumulated state.

**Fig. 5:**
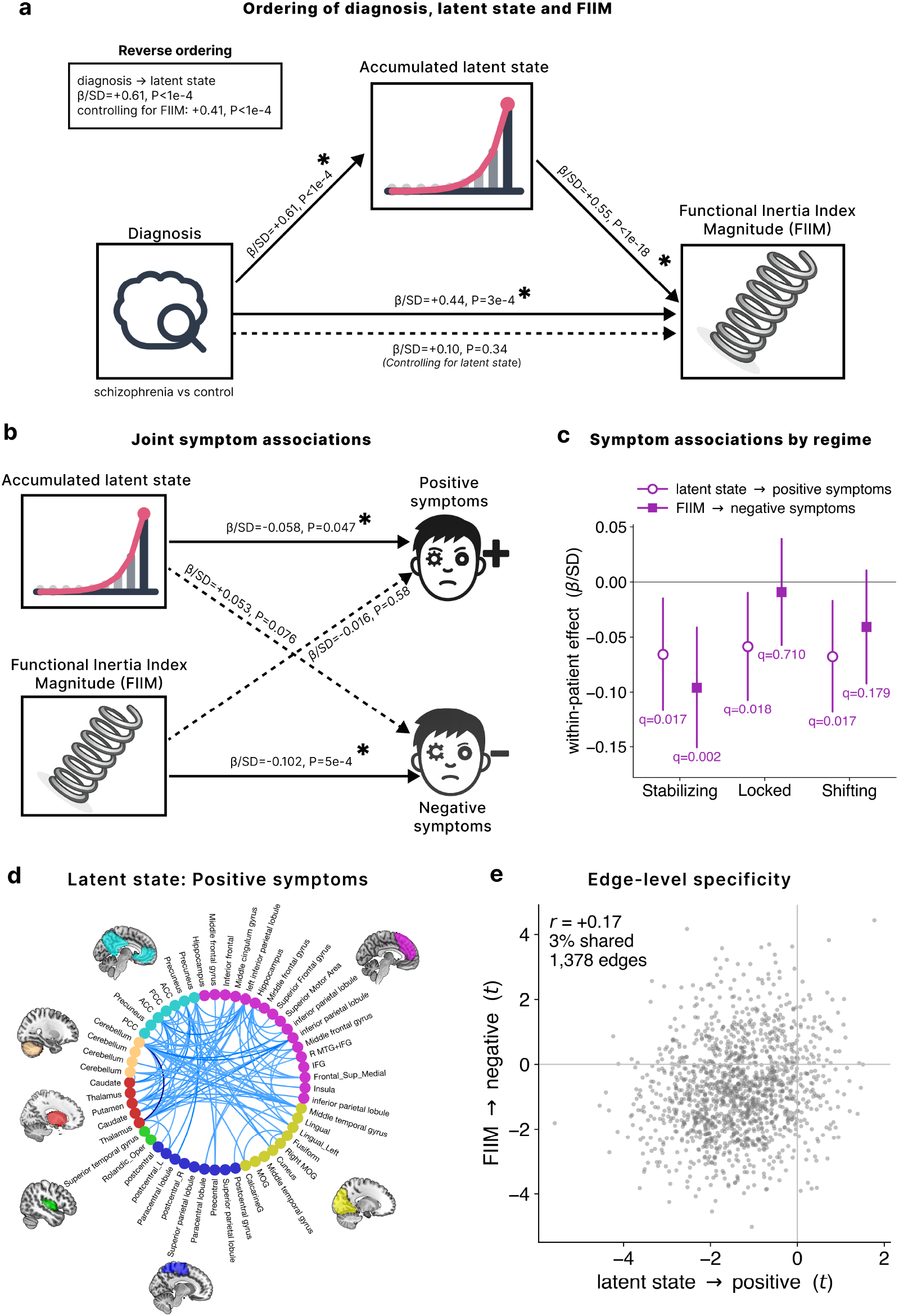
Hierarchy and symptom specificity of the accumulated latent state and functional inertia. **(a)** Ordering of diagnosis, accumulated latent state, and whole-brain functional inertia index magnitude (FIIM). Diagnosis predicts the latent state, the latent state predicts FIIM, and the diagnosis-to-FIIM association disappears once the latent state is included. The reverse ordering does not hold; diagnosis still predicts the latent state when FIIM is included in the model, establishing a specific hierarchical order. Solid arrows are significant paths; dashed arrows are not**. (b)** Latent state and FIIM entered together in a single model within patients. The latent state is associated with positive symptoms, and FIIM with negative symptoms, and neither crossed path is significant. **(c)** The same two associations were computed within each inertia regime separately. The FIIM effect is confined to the stabilizing regime, whereas the latent state effect is the same in all three, so only the FII is regime-dependent. **(d)** Edge-wise association between the accumulated latent state and positive symptom severity, drawn as in Fig. 4b. **(e)** The latent state to positive symptom map plotted against the FIIM to negative symptom map, one point per interregional pair. The two share low variance (3%), so the dissociation in (b) is spatial as well as statistical. Together, the latent state sets where the system sits and FIIM how fast it moves within that context, and the two reach different symptoms through largely different circuits.

The two also reach different symptoms (Fig. 5b). Entered together within patients, the accumulated latent state was associated with positive symptom severity (β/SD = −0.058, *P* = 0.047) and whole-brain FIIM with negative symptom severity (*β*/*SD* = −0.102, *P* = 5 × 10⁻⁴), while neither crossed path approached significance. Computing the same two associations separately within each regime shows that they are different kinds of quantity (Fig. 5c). The FIIM association with negative symptoms was confined to the stabilizing regime (*q* = 0.002) and absent in the other two, whereas the latent-state association with positive symptoms was essentially identical in all three (*q* = 0.017, 0.018 and 0.017). The functional inertia is regime-dependent, while the accumulated latent is not.

The dissociation is spatial as well as statistical. The accumulated latent state was associated with positive symptom severity in 86 of the 1,378 region pairs, every one of them in the same direction (Fig. 5d). Plotted against the corresponding map for FIIM and negative symptoms (Fig. 5e: left), the two share only 3% of their variance (*r* = +0.17; Fig. 5e), so they are different maps rather than one map relabeled. Where the system has settled (the accumulated latent state) and how fast it is still moving (functional inertia) are therefore ordered, not interchangeable, and they reach different symptom dimensions through largely different circuits.

### 3.5. Functional inertia is robust to kernel, observation model, sample, scan time, and head motion

The preceding results rest on a set of choices: how much of the past the latent state retains, what is accumulated, and which participants are included. We varied each in turn. Accumulation is what makes the index clinically informative (Fig. 6a). With accumulation removed, neither the negative symptom association in patients nor the cognitive association in controls survived. Both appeared when the past was retained, and the two behaved differently thereafter: the symptom association peaked at a finite memory and fell away as the kernel approached unbounded accumulation, whereas the cognitive association rose monotonically. The same pattern held at the edge level (Fig. 6b), where the number of region pairs surviving correction rose from 41 without accumulation to 92 at 100 s before falling to 53 under the uniform kernel for negative symptoms, and rose from 0 to 20 for reasoning and problem solving. The effects are therefore not an artefact of accumulating, and they are mostly strongest over a bounded window of recent history rather than a uniform kernel across the entire scan.

**Fig. 6:**
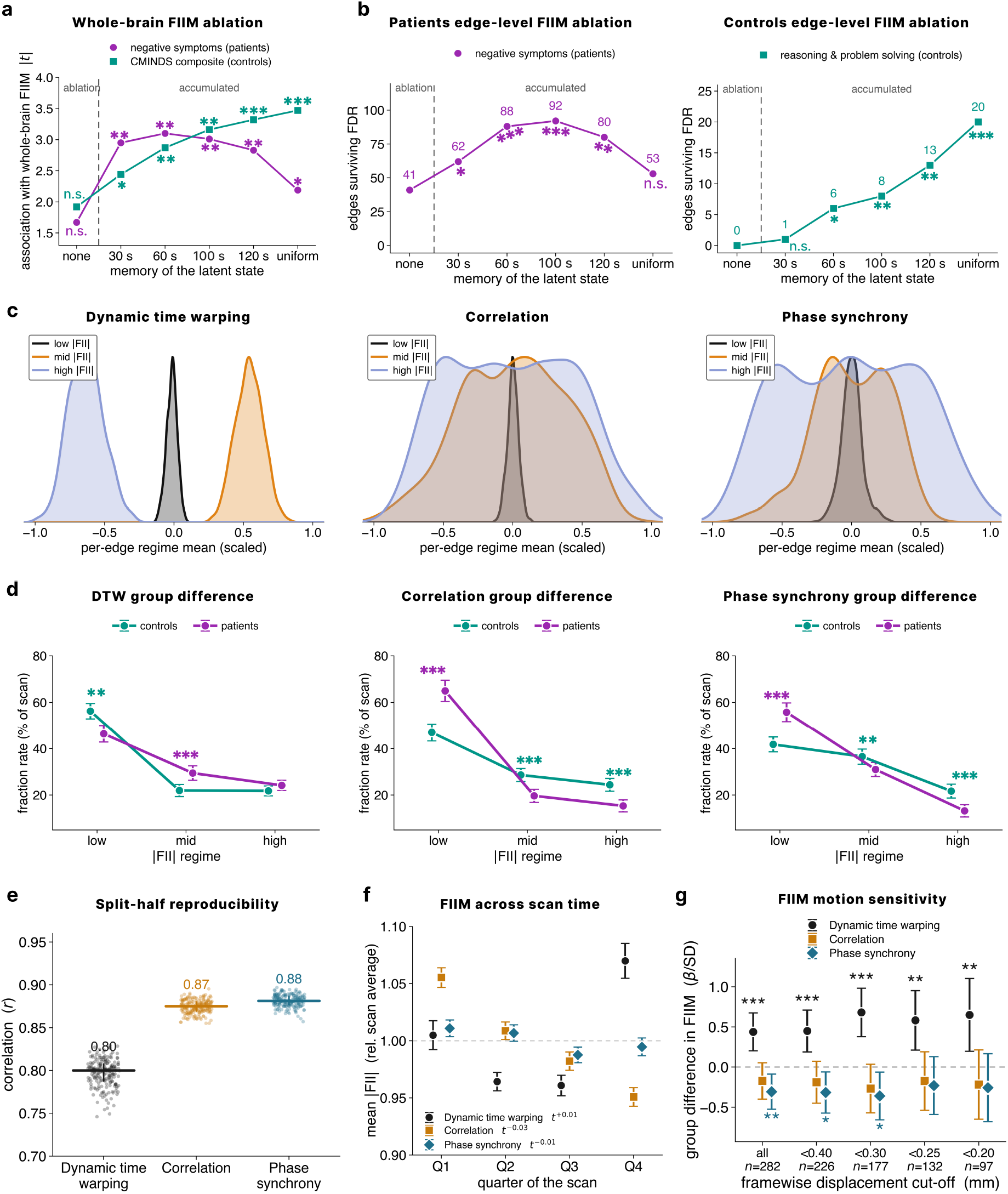
Robustness of functional inertia to kernel choice, observation model, sample, scan time, and head motion. **(a)** Subject-level associations with whole-brain FIIM as a function of how much of the past the latent state retains: no accumulation, an exponentially weighted kernel at τ = 30, 60, 100 s (the chosen kernel), and 120 s, and a uniform cumulative kernel. Negative symptoms in schizophrenia (purple) and the CMINDS composite in controls (teal) are both non-significant without accumulation. The symptom effect stays robust across all memory parameters, while the cognitive effect rises monotonically. Asterisks are the model P values. **(b)** The same ablation at the edge level, as the number of interregional pairs surviving FDR, for negative symptoms within patients (left) and reasoning and problem solving within controls (right). Asterisks are McNemar tests of each kernel against the ablation on the sets of surviving pairs. Accumulation is what makes the index clinically informative, and the effects are strongest over a finite window of the past rather than the entire scan (uniform). **(c)** Distribution of the per-edge regime means under three unrelated observations of the same data: dynamic time warping (reported observation), sliding window correlation, and phase synchrony. All three yield a three-level inertia structure, but only under DTW are the regimes cleanly single-signed and non-overlapping; under the other two they differ in magnitude alone and straddle zero. Here, regimes are therefore ranked low, mid, and high absolute FII within each model rather than named. **(d)** Fraction rate in each regime for controls and patients, under the same three observation models. The two groups separate in all three, and in each case the profiles cross, so the group difference is a redistribution of time across regimes rather than an overall shift. Asterisks are colored for whichever group is higher. **(e)** Reproducibility across participants. The sample is split at random into two halves, stratified by diagnosis and site, and the halves’ mean edge-wise FIIM maps are correlated. Each point is one of 200 splits, with the median and interquartile range. Across all three observation models, the mean correlation is above 0.8, indicating high reproducibility within the dataset. **(f)** Mean absolute FII by quarter of the scan, expressed relative to each model’s scan average, with the fitted scaling exponent beside each legend entry. A uniform cumulative kernel would force a 1/t decline; none of the three declines, and all stay within 7% of their own average. **(g)** Group difference in whole-brain FIIM under progressively stricter framewise-displacement exclusion. Patients move more than controls (mean 0.34 against 0.25 mm), yet the DTW effect is undiminished among the stillest participants. Framewise displacement is a covariate in every model throughout. Teal is controls and purple is schizophrenia; error bars are 95% confidence intervals; *** P < 0.001, ** P < 0.01, * P < 0.05.

The three-regime architecture does not depend on what is accumulated (Fig. 6c–d). Applying the identical pipeline to sliding window Pearson correlation and to phase synchrony, two observations unrelated to dynamic time warping, again produced three regimes in each, and in all three models the two groups (controls and schizophrenia) redistributed their time across those regimes. What does not transfer is the regime geometry: only under dynamic time warping are the outer regimes single-signed, so that the direction of reconfiguration is easily interpretable. Under the other two, the regimes differ in magnitude alone and all straddle zero.

Reproducibility was assessed by splitting the participants at random into two halves and correlating each half’s mean edge-level FIIM map, giving *r* = 0.80 under dynamic time warping and 0.87 and 0.88 under the other two models, across 200 splits (Fig. 6e). The index does not decay as the scan proceeds (Fig. 6f). A uniform cumulative kernel would force the FIIM to shrink as 1/*t*; instead, the EWMA kernel stayed within 7% of its own scan average in every quarter under all three observation models, with fitted exponents of +0.01, −0.03 and −0.01. Finally, patients moved more than controls (mean framewise displacement 0.34 against 0.25 mm), raising the possibility that the group difference follows from data quality rather than from inertia. Refitting under progressively stricter motion exclusion left it undiminished (Fig. 6g): the group difference in whole-brain FIIM was β/SD = +0.44 in the full sample and +0.65 among the 97 stillest participants, remaining significant at every threshold.

## 4. DISCUSSION

Large-scale brain dynamics are usually characterized by the configurations the brain occupies from moment to moment^43,44^. Functional inertia asks instead how readily the system departs from the configurations it has already accumulated, formalizing history dependence^45^ as the rate of change of an accumulated latent state. Instantiated on resting-state fMRI, this quantity is expressed at three levels: as recurrent regimes of moment-to-moment reconfiguration, as a subject-level magnitude, and as a distributed circuit-level pattern, with the accumulated state standing upstream of the rate at which it changes. A similar architecture emerges whether what is accumulated is dynamic time warping distance, sliding window Pearson correlation, or phase synchrony, although the clinical associations presented here are from dynamic time warping. The model is therefore representation-agnostic in the sense that it is well posed and productive on any time-resolved observation of brain data, not that every instantiation returns identical findings.

A system with varying reconfiguration of states cannot evolve arbitrarily^6^. It must pass through periods in which the accumulated configuration barely changes (locked), periods in which it moves toward lower interregional disparity (stabilizing), and periods in which it moves away (shifting). Clustering the functional inertia index recovers these three regimes. In the two outer regimes, every one of the 1,378 region pairs carries the same sign, so the whole brain moves in one direction at those moments, while the third sits at zero and straddles it. This matters for what the regimes are. They are not spatial states or connectivity configurations of the kind returned by conventional dynamic connectivity analyses^1,37^, and they are not a relabeling of them: the index shares no more variance with dFNC than chance allows at any window length (Figure 3g). They are modes of movement through state space under accumulated history, distinguished by the direction and the amount of reconfiguration rather than by which regions are coupled.

Schizophrenia alters how the regimes are occupied, but not in the direction a first reading suggests. Patients spend more of the scan in the stabilizing regime and dwell there longer, while controls occupy the locked regime more. Taken at face value, this would make patients the more “stable” group, which sits awkwardly beside several literature reports of reduced dynamism/connectivity in schizophrenia^3,37,46^. Inertia is contextual, and that resolves the tension. Resistance to reconfiguration preserves a coherent configuration only when the system is already in one^16^. Patients are not in one: their accumulated interregional disparity is elevated at 891 of the 1,378 region pairs and within every regime (Fig. 3b and c), including the locked one, so the displacement is a property of where they sit rather than of which regime they happen to be in. Stabilizing dynamics expressed from that starting point are convergence within a narrower dynamic range, not restoration of a well-organized configuration, and regime-conditioned comparisons confirm that no patient regime reaches the latent context occupied by controls at rest (Fig. 3c). This holds across regimes as well, with patients in the stabilizing regime remaining above controls in the locked regime (Supplementary Table 1). What distinguishes schizophrenia here is therefore a selective redistribution of time across regimes on a displaced baseline, rather than a uniform disruption of inertial dynamics.

Negative symptom severity in patients tracked inertial regime engagement. Patients as a group occupied the locked regime less than controls, yet among patients it was those who spent more of the scan locked who were the most symptomatic, while greater engagement of the shifting regime, both in time occupied and in how long each visit lasted, associated with lower symptom severity (Figure 3e). The axis separating patients from controls is therefore not the axis grading severity within them. A between-group difference and a within-group association can point in opposite directions in the same sample^47^, and we observe the same reversal in this cohort with the same DTW observation in an independent study^23^. Read within patients, what tracks severity is not how much the brain moves in total but how readily it leaves a configuration it has settled into and how long it sustains the departure. That framing is consistent with the clinical construct, in which avolition and diminished expression reflect failure to initiate and sustain a change of state rather than generalized cognitive impairment^48,49^. The association is not reducible to either of the two obvious alternatives (Fig. 3f). Adding the whole-brain accumulated latent state as a covariate left all negative symptom effects intact and slightly strengthened, so what tracks severity is the pattern of inertial regime engagement itself and not simply where the accumulated configuration sits. Adding chlorpromazine-equivalent dose, on the subset of patients for whom it was recorded, changed the coefficients by less than 0.01 and left all effects significant, so it is not an artefact of antipsychotic exposure either.

Averaged over the scan, functional inertia also behaves as a stable subject-level property, and its consequences depend on what is being held onto. In controls, whose accumulated configuration is coherent, stronger inertia was associated with better performance on the cognitive composite, on visual learning and on reasoning and problem solving (Fig. 4a). In patients whose accumulated configuration is displaced, the same measure ran the other way, with stronger inertia associated with more severe negative symptoms. Holding a well-organized configuration is useful, whereas holding a displaced one entrenches it. Hence, the reason why the measure carries opposite consequences in the two groups. The variability literature has long found that neither more nor less temporal variability is uniformly advantageous^50^, and what determines the sign here is the configuration from which the resistance operates.

The same logic applies within the brain across brain region pairs (Fig. 4b). Of the 92 region pairs tracking negative symptom severity, 77 showed stronger inertia with greater severity, and these were concentrated in cerebellar^51^, subcortical^52^, and default-mode^53^ couplings. The remaining 15 ran the other way and were largely sensory and sensorimotor, where stable coordination between regions is what the pathway requires^54^. Our results suggest that rigidity in integrative hubs keeps the system in a configuration it needs to leave, while the same resistance in sensory pathways supports the coordination those pathways depend on. This also speaks to a longstanding split in the literature, where some studies report reduced flexibility and over-stability in schizophrenia^3,37^, and others excessive variability, particularly in sensory systems^55,56^. The two need not be competing accounts. A single constraint distributed unevenly across circuits produces rigidity where high inertia entrenches a displaced configuration and volatility where low inertia degrades coordination that depends on stability.

These two quantities, where the accumulated configuration sits and how fast it changes, are derived from the same trajectory, and the relationship between them is ordered rather than symmetric (Fig. 5a). Diagnosis predicted the accumulated state, the accumulated state predicted inertial magnitude, and the association between diagnosis and inertial magnitude vanished once the accumulated state was in the model. The reverse did not hold. Patients therefore do not reconfigure faster for some independent reason. They reconfigure faster because their accumulated configuration has drifted further, which makes the elevated inertia a downstream consequence of that drift rather than a separate abnormality. The two also reach different symptoms (Fig. 5b). Entered together within patients, the accumulated latent state was associated with positive symptom severity and inertial magnitude with negative symptom severity, and the dissociation is spatial as well as statistical, since the two edge-level maps share almost none of their variance (Fig. 5c). They differ in kind as well as in consequence: the inertia effect was confined to a single regime, whereas the accumulated-state effect was the same in all three inertial regimes, so one is a property of the moment and the other a property of each subject.

Several features of these results argue that they are not artefacts of the pipeline. The accumulation step is doing the work: with it removed, neither the symptom nor the cognitive association survived (Fig. 6a and b). The effects were also strongest over a bounded window of recent history and weakened as the kernel approached unbounded accumulation. The three-regime architecture itself does not depend on what is accumulated, appearing under sliding window Pearson correlation and phase synchrony as well as dynamic time warping, with the groups redistributing their time across regimes in every case (Fig. 6c and d). What does not transfer is the regime geometry, since only under DTW are the outer regimes single-signed and the direction of reconfiguration therefore easily interpretable. The edge-level map of the inertial magnitude showed strong reproducibility across 200 random half-splits of the participants (Fig. 6e). Finally, the inertia magnitude stayed robust across scanner length and head motion (Fig. 6f and g). Taken together, this robustness makes functional inertia more likely to be a real property of brain dynamics than an artefact of the pipeline.

Several considerations bound what these results can support. The design is cross-sectional, and the associations are correlational, so the ordering we report is statistical and not a demonstration of causal direction. No perturbation was applied, and no resistance was measured directly, so functional inertia names what is measured: the amount by which an accumulated configuration changes per unit time, rather than a mechanical force opposing change. The clinical findings are specific to the dynamic time warping instantiation: the three-regime architecture and the group differences appear under correlation and phase synchrony as well, but the symptom and cognitive associations do not, so representation-agnosticism holds for the model and not for its findings. Finally, the analyses use the input-free, zero-order case of the formulation. Task-evoked designs, longitudinal data, and context-modulated accumulation kernels would extend the same model rather than alter it and would be needed to say whether the constraint behaves as it does here outside the resting state.

Large-scale brain dynamics have a dimension that is not captured by asking which configurations occur. That dimension is how much the configuration, in an accumulated context latent state, shapes what happens next. Functional inertia makes this measurable, and resolves brain activity into recurrent regimes, separates patients from controls at both the regime and whole-brain levels, and distributes across circuits in a way that accommodates both the rigidity and variability reported in schizophrenia. Its consequences are not fixed but depend on the configuration it operates from. Functional inertia therefore exposes a layer of brain organization that connectivity analyses leave unmeasured, and one that proves informative about both cognition and clinical phenotype.

## 5. ACKNOWLEDGMENT

This work was supported by the National Institutes of Health (NIH) grant (R01MH123610) and the National Science Foundation (NSF) grant #2112455.

## 6. DECLARATION OF COMPETING INTEREST

None.

## 7. DATA & CODE AVAILABILITY STATEMENT

The codes for all our analyses in MATLAB can be accessed through <u>GitHub</u>. The data was not collected by us and was provided in a deidentified manner. The IRB will not allow the sharing of data or individual derivatives, as a data reuse agreement was not signed by the subjects during the original acquisition.

## Notes

### Competing Interest Statement

The authors have declared no competing interest.

### Summary of Updates

Updates after major revisions from Translational Psychiatry. Adding more analysis and changing the kernel applied in the main. manuscript

