## supplementary material for "Functional inertia reveals history-dependent organization of large-scale brain dynamics"

### Supplementary Methods

#### DTW-based observation trajectory for the ISSM

In this work, dynamic time warping (DTW) is used to define the observed trajectory within the inertial state-space model (ISSM), providing a time-resolved measure of deviation between evolving interregional BOLD amplitude profiles. DTW serves as an observational instantiation of the ISSM, supplying the displacement-like signal from which latent state evolution and functional inertia are subsequently derived.

For each subject, DTW was applied to pairs of post-equilibrium BOLD amplitude time series $x(t)$ and $y(t)$, each of length $N$. DTW distances were computed using a Sakoe–Chiba constraint window with half-width $w$[1], selected to match the exponentially weighted moving average (EWMA) kernel, 100s/2 = 50 s. This constraint limits temporal alignment to physiologically plausible lags while preserving sensitivity to low-frequency fluctuations characteristic of resting-state dynamics[2].

The local cost function was defined as

$$\lambda_\gamma(x_i, y_j) = \left|x_i - y_j\right|^\gamma, \tag{S1}$$

where $\gamma = 1.5$. This exponent was chosen based on prior work demonstrating optimal test–retest reliability and sensitivity to clinical relevance using this value[3].

The cumulative cost matrix $CM_{(i,j)}$ for indices satisfying $\mid i - j \mid \leq w$was constructed recursively as

$$CM_{(i,j)\epsilon\{|i-j|\,\leq\, w\}} = \lambda_\gamma(x_i, y_j) + min\begin{cases} CM_{DTW(i-1,j-1)} \\ CM_{DTW(i-1,j)} \\ CM_{DTW(i,j-1)} \end{cases}. \tag{S2}$$

After completion of the cumulative cost matrix, the optimal warping path $\left(\varphi_x(\tau), \varphi_y(\tau)\right)$, with length $|\tau| \geq N$, was obtained by standard backtracking. From this path, a time-resolved DTW distance was defined as

$$D_{tr}(\tau) = \left|x\left(\varphi_x(\tau)\right) - y\left(\varphi_y(\tau)\right)\right|^\gamma. \tag{S3}$$

To obtain a uniformly sampled distance trajectory, $D_{\mathrm{tr}}(\tau)$was resampled at integer time indices $i = 1, \ldots, N$ using a piecewise cubic Hermite interpolating polynomial (PCHIP), yielding $D_{\mathrm{tr}}^{\mathrm{pchip}}(i)$ [3].

Finally, for each time point $t \geq t_a$, where $t_a$ denotes the end of the anchor baseline window, the cumulative trajectory was defined as the running cumulative average

$$c(t) = \frac{1}{t}\sum_{i=1}^{t} D_{tr}^{pchip}(i). \tag{S4}$$

This cumulative DTW trajectory constitutes the observed signal $y(t)$ in the inertial state-space model. Latent state evolution and functional inertia measures were then derived from this trajectory as described in the main Methods.

#### Estimation of the FII using DTW

For each cumulative DTW trajectory $c(t)$, the FII was estimated as the temporal derivative of the latent trajectory. Because direct differencing amplifies high-frequency noise, temporal smoothing was applied before differentiation to obtain a robust estimate of moment-to-moment reconfiguration. Specifically, a Gaussian filter was applied to each cumulative trajectory to obtain a smoothed trajectory $\tilde{c}(t)$. The filter bandwidth was selected in a data-driven manner based on the frequency content of $c(t)$. The mean power spectral density (PSD) of the cumulative trajectories was computed across subjects, and the frequency $f_p$ at the elbow of the power log of the PSD was identified. This frequency was used as the –3 dB (half-power) cutoff of the Gaussian filter.

The frequency response of the Gaussian filter is given by

$$|H(f)|^2 = e^{-4(\pi f \sigma_t)^2}, \quad \text{(S5)}$$

where $\sigma_t$denotes the temporal standard deviation of the filter. Setting $|\, H(f_p)\,|^2 = 1/2$ yields

$$\sigma_t = \sqrt{\frac{\ln 2}{4\pi^2 f_p^2}}. \quad \text{(S6)}$$

The corresponding full width at half maximum (FWHM) in the time domain was computed as

$$FWHM = 2\sqrt{2\ln 2}\,\sigma_t\,, \quad \text{(S7)}$$

and converted to samples based on the repetition time (TR) of the acquisition. The smoothed cumulative trajectory $\tilde{c}(t)$ was then differentiated using first differences,

$$\dot{c}(t) = \tilde{c}(t) - \tilde{c}(t-1)\,, \quad \text{(S8)}$$

to obtain the FII. For the present dataset (TR = 2 s), this procedure yielded a Gaussian filter with FWHM of 4 samples (Supplementary Fig. 1).

Because DTW distances are accumulated over time, the sign of the discrete derivative reflects whether the cumulative deformation cost is increasing or decreasing, corresponding to divergence from or convergence toward the running configuration, respectively. For interpretability, the discrete derivative was multiplied by −1 so that positive values correspond to convergence and negative values to divergence, without altering the magnitude of the effect. Temporal smoothing suppresses high-frequency noise while preserving meaningful reconfiguration events, yielding a stable estimate of functional inertia.

#### Cross-regime comparison of latent-state context

Regime-conditioned latent-state magnitude was computed by averaging the latent trajectory across all time points assigned to each regime, giving one value per regime per participant. These were compared between groups across all nine combinations of patient and control regimes using a repeated-measures ANCOVA adjusted for age, sex, scanning site, and head motion, with false discovery rate correction across comparisons. Latent-state magnitude in schizophrenia exceeded that of controls for every pairing (Supplementary Table 1). Patients in the stabilizing regime remained above controls in the locked regime, so directed convergence in schizophrenia does not reach the latent-state context of stable configurations in controls.

Supplementary Figure 1 | Spectral knee selection across the subject-averaged PSD

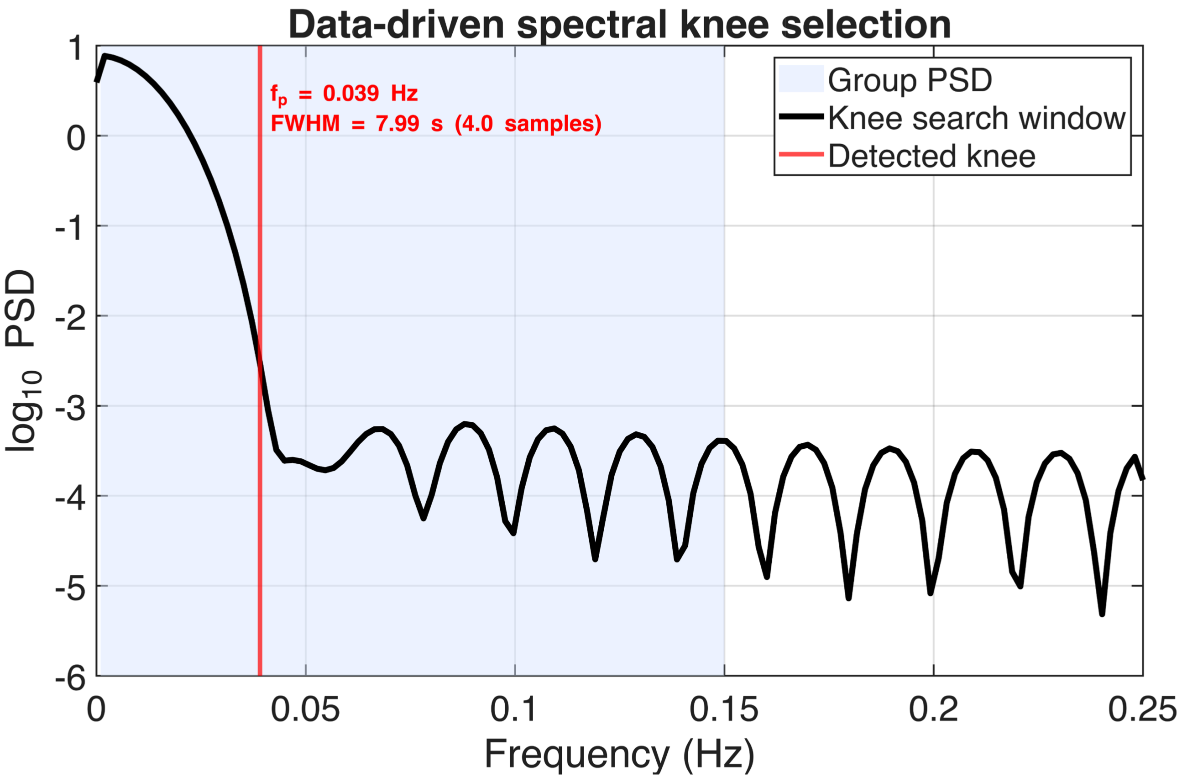


*Supplementary Figure 1 Spectral knee selection across subject-averaged PSD of the cumulative DTW trajectory. The power spectral density (PSD) of the cumulative DTW trajectory was averaged across all subjects and brain region pairs and plotted on a logarithmic scale. The shaded band denotes the frequency window used for knee detection. The vertical line marks the detected spectral knee frequency $f_p$, representing the transition from signal-dominated low-frequency structure to higher-frequency fluctuations. This knee was used to define the –3 dB cutoff of a Gaussian smoothing filter, yielding a data-driven full width at half maximum (FWHM) used in subsequent temporal smoothing.*

**Supplementary Figure 2 | Determination of the number of functional inertia regimes**

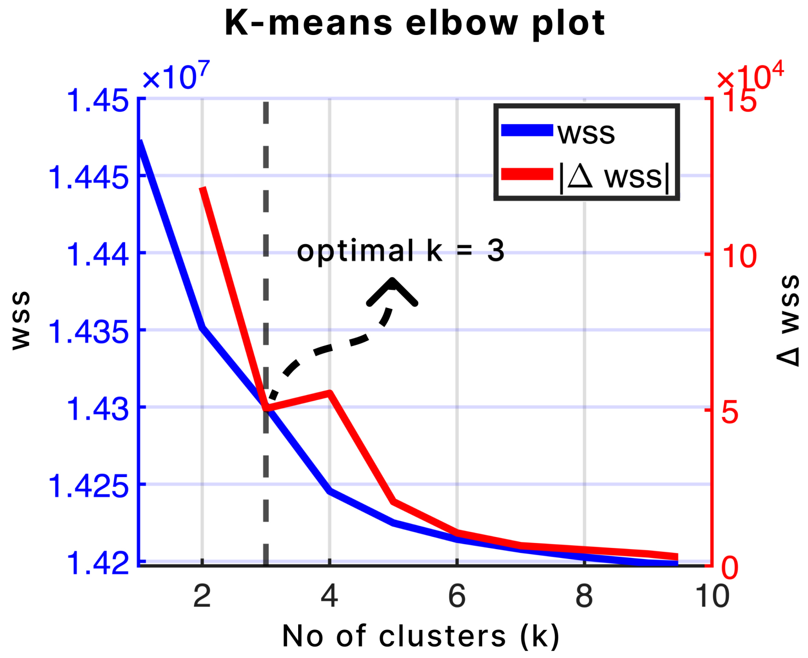


*Supplementary Figure 2 Elbow analysis for k-means clustering of the Functional Inertia Index (FII) time series concatenated across subjects and interregional pairs. The within-cluster sum of squares (WSS; blue) decreases monotonically with increasing cluster number, while the change in WSS between successive solutions (|ΔWSS|; red) exhibits a clear inflection at k = 3, indicating diminishing returns beyond this point. This pattern supports the selection of three clusters, corresponding to the locked, stabilizing, and shifting functional inertia regimes used*

*Supplementary Table 1 | Cross-group, cross-regime comparisons of latent-state magnitude*

| **Schizophrenia Regime** | **Control Regime** | **Δ (Schizo. – Control)** | **95% CI** | **p-value** | **pFDR** |
|---|---|---|---|---|---|
| Stabilizing | Locked | 0.01608 | [0.01073, 0.02143] | $1.1 \times 10^{-8}$ | $1.1 \times 10^{-8}$ |
| Stabilizing | Stabilizing | 0.01586 | [0.01046, 0.02127] | $9.0 \times 10^{-9}$ | $1.35 \times 10^{-8}$ |
| Stabilizing | Shifting | 0.01453 | [0.00921, 0.01985] | $2.4 \times 10^{-7}$ | $2.9 \times 10^{-7}$ |
| Locked | Locked | 0.01482 | [0.00950, 0.02014] | $4.8 \times 10^{-8}$ | $4.8 \times 10^{-8}$ |
| Locked | Stabilizing | 0.01455 | [0.00918, 0.01992] | $6.1 \times 10^{-8}$ | $6.8 \times 10^{-8}$ |
| Locked | Shifting | 0.01322 | [0.00794, 0.01850] | $1.7 \times 10^{-6}$ | $2.1 \times 10^{-6}$ |
| Shifting | Locked | 0.01716 | [0.01183, 0.02249] | $3.5 \times 10^{-9}$ | $5.4 \times 10^{-9}$ |
| Shifting | Stabilizing | 0.01583 | [0.01058, 0.02109] | $8.9 \times 10^{-9}$ | $1.35 \times 10^{-8}$ |
| Shifting | Shifting | 0.01556 | [0.01029, 0.02083] | $7.2 \times 10^{-9}$ | $1.35 \times 10^{-8}$ |

*Table 1 Cross-group comparisons of regime-conditioned latent-state magnitude between schizophrenia and control participants across all combinations of functional inertia regimes. Differences represent estimated marginal mean contrasts (Schizophrenia minus Control) derived from the repeated-measures ANCOVA model, adjusted for age, sex, scanning site, and head motion. Positive values indicate higher interregional disparity in schizophrenia. False discovery rate (FDR) correction was applied across all cross-regime comparisons.*
